# Biogeochemical function of slicks in coastal surface waters of the Baltic Sea

**DOI:** 10.1101/2025.06.25.660598

**Authors:** Carolin Peter, Helge-Ansgar Giebel, Bingli Clark Chai, Tassiana S. G. Serafim, Carola Lehners, Oliver Wurl, Helena Osterholz, Janina Rahlff

## Abstract

The sea-surface microlayer (SML) is a crucial ocean-atmosphere interface involved in gas exchange and nutrient cycling. Slicks, i.e., viscous surface layers, common in coastal regions serve as microbial hotspots. We studied microbial abundance, surfactants, dissolved organic carbon (DOC), and net community production (NCP) of O_2_ in slick and non-slick SMLs and underlying water (ULW) in the coastal Baltic Sea. Slicks often showed higher surfactant levels compared to the ULW. Microbial respiration often exceeded production, resulting in net O_2_ consumption, although some ULW sites exhibited net O_2_ production. The SML was enriched with pico- and nanophytoplankton, with cyanobacteria being negatively correlated with total dissolved nitrogen. In contrast, microphytoplankton accumulated in the ULW, indicating niche separation with depth. Microscopy revealed ciliates and juvenile sporophytes dominating a slick’s >100 µm fraction. In eutrophic coastal systems, slicks influence plankton communities and O_2_ dynamics, supporting their role in surface biogeochemical cycling and climate-driven changes.

## Introduction

The sea-surface microlayer (SML) constitutes a 1-mm thick boundary layer between ocean and atmosphere and covers 70% of Earth’s surface. In its interfacial position, the SML has a critical role in gas exchange, nutrient cycling, and the transfer of organic and inorganic materials (Engel et al. 2017). In addition, the thin but highly reactive SML is enriched with various substances, including surface-activate substances (surfactants), dissolved organic carbon (DOC), and microbes, which influence SML dynamics (Hardy 1982). The enrichment of surfactants, e.g. derived from biogenic production (Ẑutić et al. 1981), is usually a prerequisite for slicks to form and persist, and the appearance of slicks in coastal regions is more common than in the open sea (Romano and Marquet 1991; Romano 1996). At low wind speed, when bacteria tend to accumulate in the SML (Rahlff et al. 2017a; Stolle et al. 2010), their accumulation can become visible at the water surface and form a viscous surface slick (Carlson 1987; Dietz and Lafond 1950). Microbial activity, especially heterotrophic activity, is often enhanced in the SML compared to the underlying water (ULW) (Obernosterer et al. 2005; Rahlff et al. 2017b; Reinthaler et al. 2008). Slicks have been described as an important habitat for various microbes (reviewed by Voskuhl and Rahlff (2022)), including fungi, bacteria and viruses (Crow et al. 1975; Rahlff et al. 2023b; Sieburth and Conover 1965; Wurl et al. 2016) but are known to support higher-level organisms of the food web, for example as nurseries (Gallardo et al. 2021; Helm 2021; Whitney et al. 2021). Microorganisms in and under the SML can metabolically influence gas exchange across the air-sea boundary (Calleja et al. 2005; Upstill-Goddard et al. 2003). When oxygen (O2) consumption, used as a proxy for metabolic activity, was measured in enclosed SML samples collected under slick-like conditions from an aquarium tank, rapid oxygen depletion was observed, and anoxia was reached within a few hours due to elevated microbial metabolic activity (Rahlff et al. 2019). The net community production (NCP) rates of O_2_ showed strong variations between -21.3 and 44.8 μmol O_2_ L^−1^ h^−1^, indicating that both respiration and primary production can be enhanced under slick-like conditions (Rahlff et al. 2019). Equivalent measurements from field SML slicks are currently lacking, as is an understanding of the relationship between microbial activity, reflected in O_2_ consumption rates, and surfactant presence in slicks. Surfactants in slicks typically reduce surface tension at the air-sea interface (Romano and Marquet 1991; Sturdy and Fischer 1966), which, in principle, could facilitate the aggregation and sinking of organic material that forms the basis of marine snow (Quigg et al. 2021). Here, we investigated the relationship between microorganisms, surfactants, DOC, and NCP rates of O_2_ in and below the SML of slicks from the natural marine environment to gain a deeper understanding of their influence on biogeochemical processes regulating O_2_ dynamics in the upper surface ocean.

## Methods

### Sampling

Slick SML (SMS), non-slick SML (SMN), slick underlying water (ULS), and non-slick underlying water (ULN) samples were collected from the Baltic Sea off the coast of Warnemünde over three days (13^th^ =Day1, 16^th^=Day2, 18^th^=Day3 June 2024, Figure 1A). SML was sampled using a glass plate sampler (Harvey and Burzell 1972), as the SML adheres to the glass due to surface tension forces. After dipping and slow withdrawal of the glass plate from the bow of a small boat, the sample was collected with a squeegee (Figure 1C) through a 100 µm mesh size plankton sieve (Figure 1D, Aquacopa, Jabel, Germany) into a 1-L brown HDPE bottle (Nalgene, Rochester, USA). The glass plate, funnel, and sampling bottles were cleaned with household bleach, ethanol, and sample water before use. For reference purposes, ULW samples were collected from 1 m depth using a syringe connected to a weighted hose. In the field, we collected data for salinity and water temperature using a portable TDS meter CO-330 (VWR, Darmstadt, Germany). Wind speed was averaged from readings on a hand-held anemometer model MS6252A (Mastech Group, Brea, CA, USA) held ∼1-2 m above the water surface. Light intensity was measured on the boat using the Galaxy Sensors phone app v.1.10.1.

**Figure 1:**
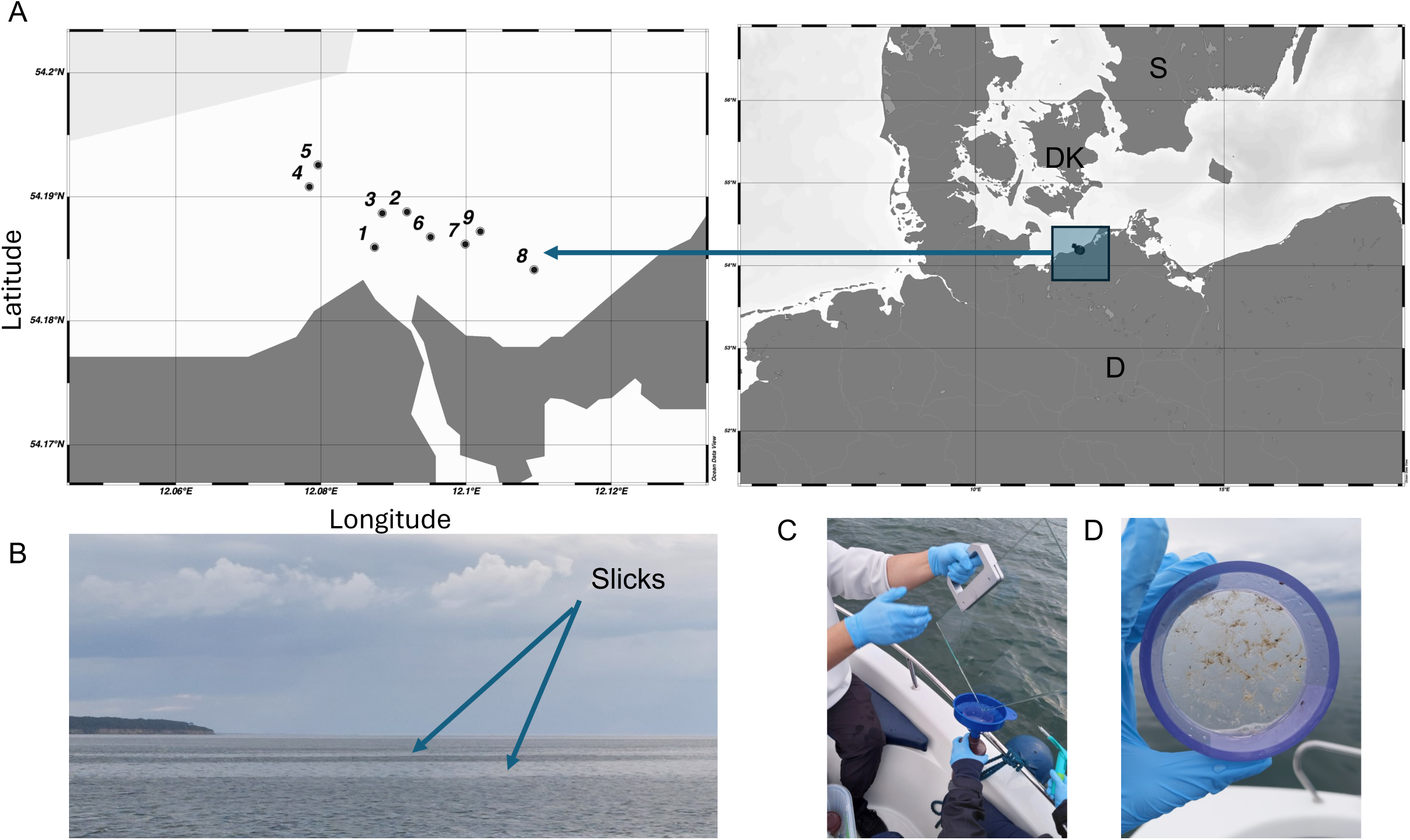
Map of sampling stations (A) in the Baltic Sea. For details corresponding to the different slick samples, see Table S1. Representative slicks in the coastal Baltic Sea emerging on 16^th^ June 2024 (B). Example of glass plate sampling method (C), and plankton sieve with 100 µm mesh filled with slick SML biomass (D). Photos by Janina Rahlff; map constructed using Ocean Data View v. 5.7.2 (Schlitzer 2022).

### Dissolved organic carbon (DOC) and total dissolved nitrogen (TDN)

For DOC and TDN analysis, 2 x 15 mL of sample water were pushed through pre-combusted (400 °C for 4 h) 47 mm diameter GF/F filters (Whatman, Buckinghamshire, UK, pore size 0.7 μm) into pre-combusted glass vials closed with acid-washed lids including PTFE-lined septa and were stored frozen until analysis at -20 °C. In some cases, only a single measurement was available (see Table S2). DOC and TDN concentrations were obtained via total organic carbon analyzer high temperature catalytic combustion (TOC-L, Shimadzu, Duisburg, Germany) calibrated with acetanilide. Expanded measurement uncertainties were 7.7 % for DOC and 8.1 % for TDN.

### Surfactant analysis

Remaining sample water (∼100 mL volume) of SML and ULW for surfactant analysis were shipped to Oliver Wurl’s lab on cool packs. The samples, with some slick samples needing dilution, were analyzed using an established voltametric technique with a hanging mercury drop electrode (Rickard et al. 2019). Surfactants were quantified using the standard addition technique. The non-ionic surfactant Triton X-100 (Sigma-Aldrich, Taufkirchen, Germany) was used as a standard, and surfactant concentration is expressed as the equivalent concentration of Triton X-100 (µg Teq L^−1^). Biogeochemical functions of slicks

### Cell count analysis

Samples were preserved with a 1% solution of glutaraldehyde (Carl Roth, Karlsruhe, Germany), kept in the dark for an hour, and then frozen at -80 °C. Slick samples, which are naturally more enriched with particles, were filtered through 50 µm CellTrics® filters (Sysmex Partec, Muenster, Germany) using gravity filtration prior measurements to avoid clogging of the device. Cell counts were determined using a BD Accuri C6 flow cytometer (Becton Dickinson Biosciences, Franklin Lakes, USA) according to Giebel et al. (2021) and Marie et al. (2000). Small, autofluorescent phototrophic eukaryotes (phytoplankton) were quantified following the gating strategy shown by John et al. (2022) with slight modifications (a threshold of 600 on the FL3 channel) and revealed populations of micro-, smaller and larger nano- (N2 and N3), and picophytoplankton, which can contain cyanoprokaryotes, but lacking a FL2-signal (Figure S2). Cytograms showed for several samples a cyanobacterial population (Cyano) derived from its FL3/FL2-signals. Prokaryotic abundance was assessed using a modified protocol by Giebel et al. (2021) and Rahlff et al. (2023a), in which the samples were stained with SYBR® Green I (10x, Invitrogen/Thermo Fisher Scientific, Carlsbad, CA, USA). Fluorescent latex beads (1 μm, Polysciences Europe, Eppelheim, Germany) served as internal controls and performance reference.

### Microscopic analysis

During encounter of a particular viscous slick at Station 4 on the 16^th^ June 2024 (Figure 1B), which featured foams, seaweeds, insects, and potentially cyanobacterial filaments (Figure S3), we collected the biomass retained on the 100 µm mesh size plankton sieve (Figure 1D) with a cleaned glass scraper for microscopic analysis. The sample remained uncooled during shipment and before fixation with acidic Lugol’s iodine solution at a final concentration of 1% and subsequent storage in the dark at 4 °C. Microscopic analysis was performed within six months using an inverted light microscope (CK X 41, Olympus, Evident, Tokyo, Japan) at 200 - 400 x magnification. For community composition analysis, particles, cells and organisms in 820 µL were enumerated.

### Oxygen consumption measurements

We used the FireSting® needle-type oxygen minisensor OXF500PT (PyroSience GmbH, Aachen Germany) to measure dissolved O_2_ of SML and ULW samples in 3-mL Exetainers^®^ vials (Labco Limited, Ceredigion, UK), which were continuously mixed by a small stirrer bar. The exetainers were submerged in a water-filled beaker during measurements. The sensor was calibrated daily before use in air-saturated water, which was achieved by bubbling tank water with ambient air for 15 minutes (100% O_2_), and by using an anoxic 0.1 M Sodium L+Ascorbate solution (0% O_2_) (Sigma-Aldrich/Merck, Darmstadt, Germany). Duplicate samples were measured either under ambient light conditions to facilitate both primary production and O_2_ consumption (GPP=gross primary production) or in exetainers wrapped with aluminum foil to measure O_2_ consumption (respiration) exclusively. O_2_ fluxes were also measured in 0.2 µm-filtered samples to account for non-biological O_2_ fluxes. However, these measurements showed unexpected variability, preventing clear conclusions about non-biological O_2_ dynamics. Due to inconsistent results and potential undetected factors affecting oxygen fluxes, we did not consider these measurements to be reliable negative controls. Temperature in the water-filled beaker into which the exetainer was submerged during measurement was continuously recorded with a TDIP15 probe (PyroSience GmbH).

### Analysis of net community production of O_2_

Data were analyzed in PyroScience Data Inspector v.1.5.3.2466 by using the salinity value recorded for the specific day (Table S1) and corrected for the temperature measured alongside, based on the salinity and temperature dependence of O_2_ solubility (Garcia and Gordon 1992). The linear respiration rate function of the software was used. A 10-minute measurement window was selected (Figure S1). On Day 1, this spanned 10 - 20 of a 30-minute recording to allow time for initial sensor stabilization. On Days 2 and 3, the full 10-minute measurement period was used. Recordings on these days began only after an initial sensor adjustment period following its injection into the sample. Data analysis was conducted using Microsoft Excel to calculate Net Community Production (NCP) from Gross Primary Production (GPP), and Respiration (R) according to the formula: NCP = GPP - R. The mean values of GPP and R from duplicate measurements were calculated, and NCP was then derived by subtracting the mean R from the mean GPP.

### Enrichment factors (EF) and statistics

The enrichment factor (EF) is used to quantify the concentration difference of a parameter between the SML and the ULW, calculated using the formula EF = CSML / CULW, where CSML is the concentration of the parameter in the SML and CULW is its concentration in the ULW (GESAMP 1995). EF values > 1 indicate enrichment in the SML, whereas EF values ≤ 1 indicate depletion. Plots, heatmaps and correlations (matrices) were performed using GraphPad Prism v.10.4.2 (Boston, Massachusetts, USA). Correlations were two-tailed, non-parametric Spearman correlations at the 95% confidence interval.

## Results

### Environmental conditions and dissolved compounds

In situ water temperatures of the Baltic Sea during sampling ranged between 15.5 and 16.9 °C, salinity between 11.2 and 12.0, and wind speeds between 1 and 5.5 m s^-1^. DOC values were slightly higher in the SML (SMS+SMN at 340 - 477 µM, compared to the ULW (318 - 426 µM in ULS+ULN (Figure 2A) indicating minor EFs of maximum 1.1. Total dissolved nitrogen (TDN) ranged from 19 to 30 µM for SML and from 15 to 26 µM in ULW (Figure 2A) leading to EFs between 0.9 and 1.3. Surfactant concentrations were also higher in SMS+SMN samples at 200.4 - 825.9 µg Teq L^-1^ than in ULS+ULN samples at 108 - 364.1 µg Teq L^-1^ (Figure 2B). Surfactant EFs on Day1 indicated depletion (0.6 and 0.7) for the two slicks sampled. Day2 and Day3 SML samples’ EFs ranged between 1.7 and 4.0 indicating medium to strong surfactant enrichments (Table S2).

**Figure 2:**
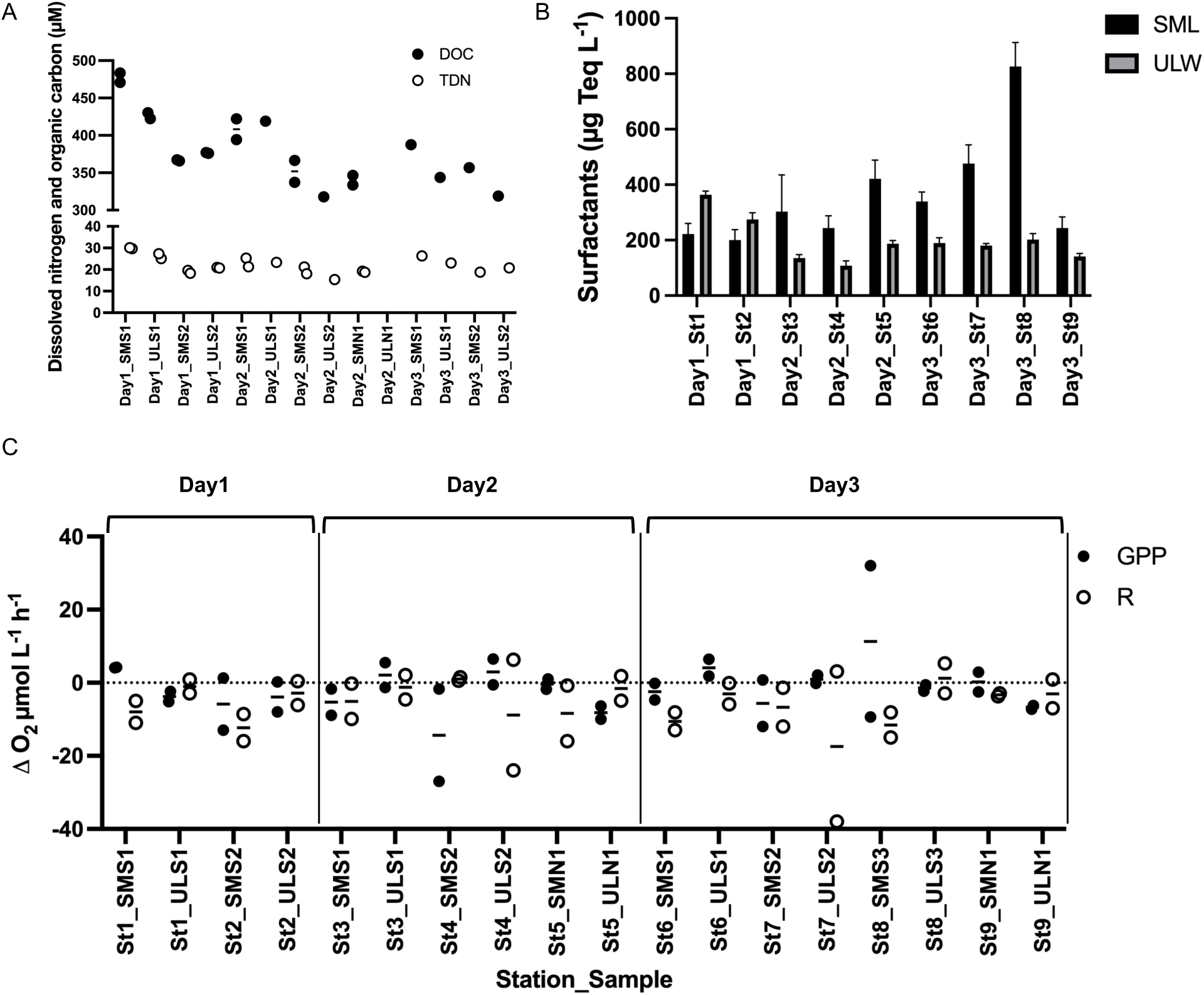
Total dissolved nitrogen and organic carbon (TDN, DOC) (A) and surfactants (B) in the surface microlayer and underlying water samples for three different days. Gross primary production (GPP) and respiration (R) expressed as measured in the samples from the delta O_2_ µmol L^-1^ h^-1^ (C).

### Microbial abundance across size classes

Microbial abundance was typically higher in SML than in ULW samples and declined with increasing cell size. Prokaryotic cell counts ranged from 3.1 - 9.2 x 10^6^ cells mL^-1^ in SMS+SMN samples and from 2.2 - 3.9 x 10^6^ cells mL^-1^ in ULS+ULN samples (Figure 3A), indicating EFs of 1.2 - 2.5. Cyanobacterial abundance ranged from 0.74 - 3.4 x 10^5^ cells mL^-1^ in SMS+SMN samples, and from 0.77 - 1.4 x 10^5^ cells mL^-1^ in ULS+ULN samples (Figure 3A). Cyanobacterial EFs ranged from 0.8 - 2.2. Smallest primary producers, the picophytoplankton ranged from 0.31 - 1.3 x 10^5^ cells mL^-1^ in SMS+SMN samples and from 3.0 - 6.2 x 10^4^ cells mL^-1^ in ULS+ULN samples with EFs from 0.8 - 2.7. Small nanophytoplankton (population N2 in Figure S2) ranged from 0.57 - 2.0 x 10^4^ cells mL^-1^ in SMS+SMN samples, from 5.2 - 7.7 x 10^3^ cells mL^-1^ in ULS+ULN samples and EFs from 0.8 - 2.3. Larger nanophytoplankton (population N3 in Figure S2) ranged from 0.03 - 2.7 x 10^4^ cells mL^-1^ in SMS+SMN samples, from 1.1 - 8.7 x 10^2^ cells mL^-1^ in ULS+ULN samples and EFs from 0.6 - 17.6. Microphytoplankton ranged from 212 to 1169 cells mL^-1^ in SMS+SMN samples, from 48 - 632 cells mL^-1^ in ULS+ULN samples, and EFs ranged from 1 - 9.7.

**Figure 3:**
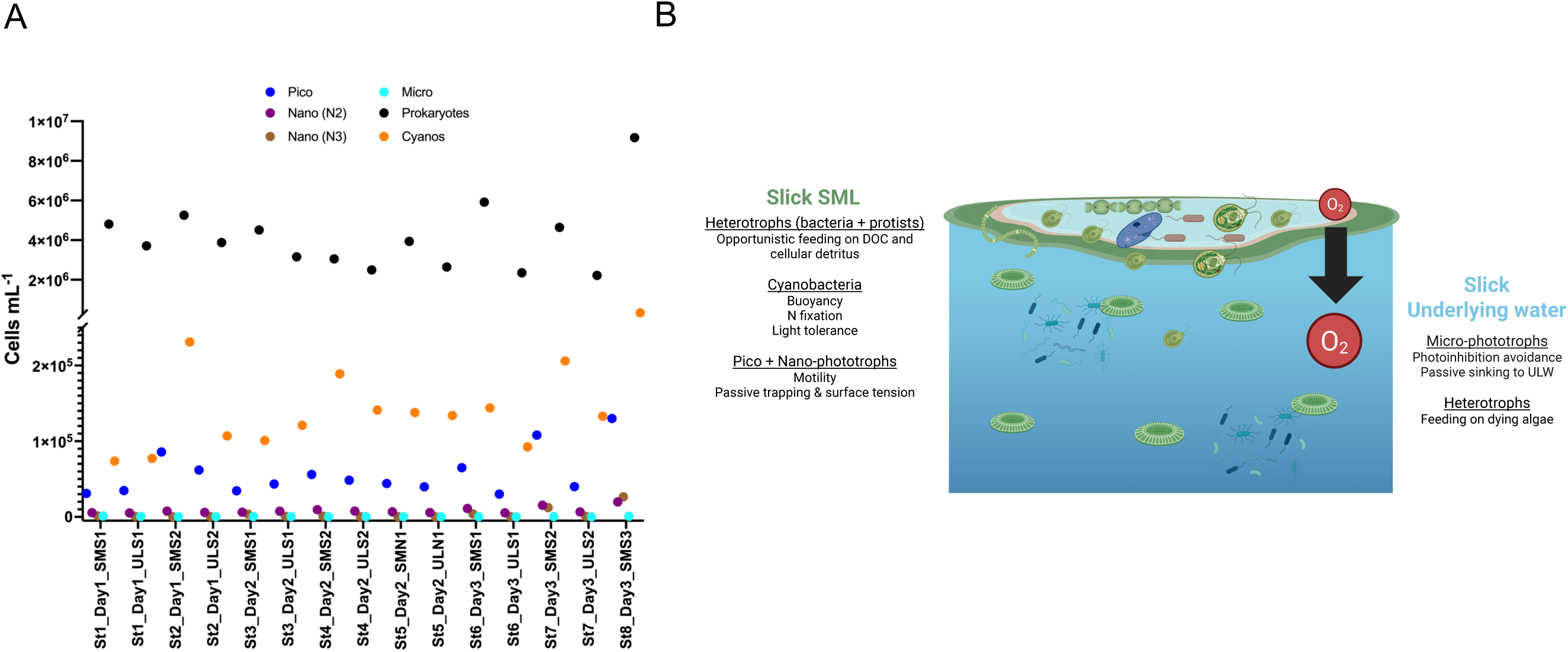
Abundances of different unicellular populations as cells mL^-1^ across three sampling days (A), concept figure of typical groups of accumulating microorganisms in and underneath slicks (B).

### Production and consumption of O_2_

Across all days, respiration (R) was consistently more variable than GPP, and in 5 out of 7 slick sites (SMS), R exceeded GPP, indicating a system where heterotrophic activity played a dominant role in O_2_ dynamics (Figure 2C). On the first day, the strongest O_2_ depletion was observed in SMS2, where Δ O_2_ of GPP dropped to -13 μmol O_2_ L^−1^ h^−1^ and Δ O_2_ of R to -16 μmol O_2_ L^−1^ h^−1^. GPP remained relatively low across all sites, with maximum values of 4.2 μmol O_2_ L^−1^ h^−1^ in SMS1. It indicates that heterotrophic processes dominated O_2_ fluxes, leading to a net O_2_ loss. On Day2, variability increased, with some sites showing NCP of O_2_, particularly in ULS1, SMN1, and ULS2. By the third day, the range of Δ O_2_ values widened further. SMS3 exhibited the highest O_2_ production, with GPP reaching up to 32 μmol O_2_ L^−1^ h^−1^ indicating favorable conditions for primary production. However, several locations (SMS1, SMS2, ULS2) continued to show O_2_ consumption, with respiration rates reaching -38 μmol O_2_ L⁻¹ h⁻¹. Overall, the slick sites tended to have the lowest Δ O_2_ values, indicating stronger heterotrophic activity and/or lower primary production (Figure 2C). Conversely, ULS and ULN sites occasionally supported NCP of O_2_, suggesting that microbial dynamics differed between the SML and deeper layers. The patchy distribution of O_2_ turnover rates aligns with the heterogeneous nature of the SML, where local environmental factors and microbial community composition likely contribute to the observed variability.

### Correlations between NCP of O_2_ and predictors

Significant positive Spearman correlations in the SML (including SMS+SMN) were observed between picophytoplankton and nanophytoplankton N2 (rho=0.93, *p*=0.002), picophytoplankton and cyanobacteria (rho=0.95, *p*=0.001), as well as nanophytoplankton N2 and cyanobacteria (rho=0.83, *p*=0.015) (Figure S4). Furthermore, a significant negative correlation was observed for cyanobacteria and TDN (rho=-0.79, *p*=0.048, Figure 4A). Prokaryotes and NCP of O_2_ show positive correlations (rho=0.76, *p*=0.037) in the SML (Figure 4B). In the ULW (including ULS+ULN), significant positive Spearman correlations were found for TDN and DOC (rho=0.94, *p*=0.017), prokaryotes and microphytoplankton (rho=0.82, *p*=0.034, Figure 4C), while significant negative correlations were found for surfactants and nanophytoplankton N2 (rho=-0.89, *p*=0.012) as well as for DOC and NCP of O_2_ (rho=-0.89, *p*=0.033). The EFs of picophytoplankton correlated significantly and positively with EFs of nanophytoplankton N2 (rho=1.00, *p*<0.001) and EFs of cyanobacteria (rho=0.86, *p*=0.024). These patterns point toward co-enrichment, likely influenced by shared physical and biological drivers. In addition, there were correlations between SML and ULW features: DOC and TDN were significantly positively correlated between both layers (rho=0.94, *p*=0.017 and rho=1.00, *p=*0.003, respectively), showing that SML and ULW are chemically interconnected. DOC in SML and nanophytoplankton N3 in the ULW were significantly positively correlated (rho=0.86, *p*=0.024). Surfactants in SML were significantly negatively correlated with prokaryotes in ULW (rho=-0.82, *p*=0.034). NCP of O_2_ in the SML was positively correlated to surfactants in ULW (rho=0.85, *p*=0.006) and negatively with nanophytoplankton N2 in the ULW (rho=-1.00, *p*<0.001). Nanophytoplankton N2 in the SML was negatively correlated with microphytoplankton in the ULW (rho=-0.86, *p*=0.024) and with prokaryotes in the ULW (rho=-0.79, *p*=0.048). Microphytoplankton in the SML was significantly positively correlated with nanophytoplankton N3 in the ULW (rho=0.88, *p*=0.015). Spearman correlation coefficients are shown in Figure 4 D-F and Table S3. A concept figure summarizes the distributions of planktonic and neustonic microbes in the slick ecosystem (Figure 3B).

**Figure 4:**
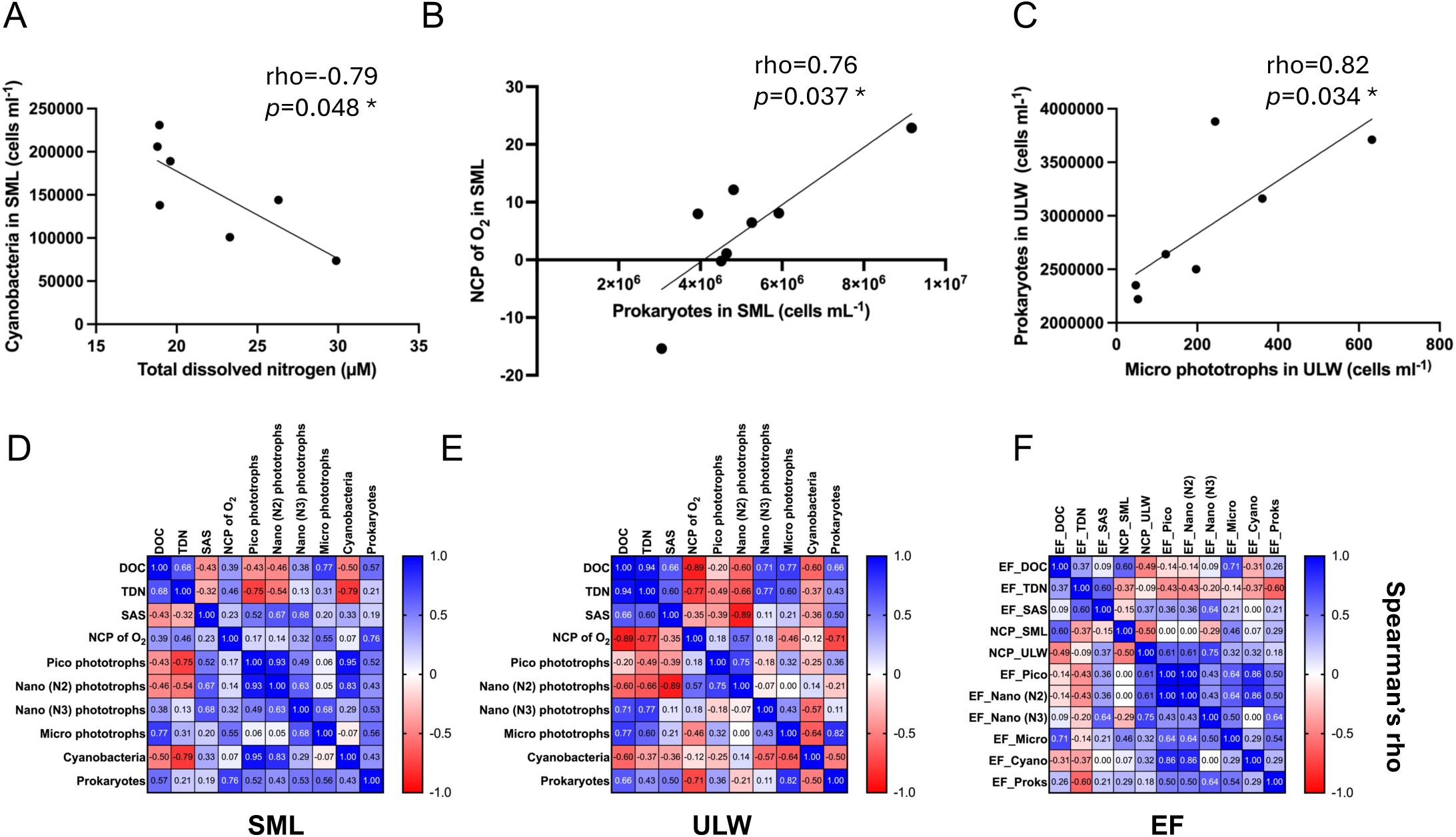
Negative relationship between cyanobacteria and total dissolved nitrogen in the SML (A), positive relationship between prokaryotes and NCP of O_2_ in the SML (B) and positive relationship between prokaryotes and microphytoplankton in the ULW (C); Correlation matrices for biological and physico-chemical parameters within the sea-surface microlayer (SML) (D), the underlying water ULW (E) and between enrichment factors (EF) (F), i.e., concentration in the SML divided by concentration in the ULW for these parameters. Spearman’s rho and *p*-values for correlations are shown in Table S3. EF= enrichment factor, DOC=Dissolved organic carbon, SAS=surface-active substances (surfactants), SML=sea-surface microlayer, TDN=Total dissolved nitrogen, ULW=underlying water

### Microscopic analysis of a slick >100 µm fraction

The microscopic sample was dominated by loosely aggregated detritus. It was comprised of microscopic particles such as pollen and dead algal cells, as well as parts of metazoans such as stellate hairs, bristles, lepidopteran scales or parts of shed exoskeletons. The autotrophic community was comprised mostly of juvenile sporophytes (10^3^ mL^-1^) and accompanied by pennate diatoms (17 cells mL^-1^, Figure 5A), different chlorophytes, dinoflagellates, and filamentous cyanobacteria (Figure 5D, Table S4). The microzooplankton was dominated by three distinct ciliate types at 7, 1, and 0.03x10^3^ cells mL^-1^ (Figure 5E). The most and least common types shared a flexible body resembling *Gastrostyla* sp. or *Uroleptus* sp. (Figure 5E), the third type resembled a contracted and armoured *Vorticella* sp. (Figure 5B). All three would have been expected to pass through the sieve based on their size (30-35 µm length x 10-12 µm width, ø=30 µm) suggesting entrapment in the detrital matrix. Copepods and cladocerans were absent from the samples. The mesozooplankton community consisted mostly of nematodes (10^3^ mL^-1^) together with water mites of which almost exclusively exoskeletons were found. It is noteworthy that the high density of two metazoan organisms, one distinctly resembled the fish parasite *Gnathia* sp. (10 mL^-1^, Figure 5C) and the other with a body structure resembling that of miniature trichopteran larvae (17 mL^-1^, Figure 5F).

**Figure 5:**
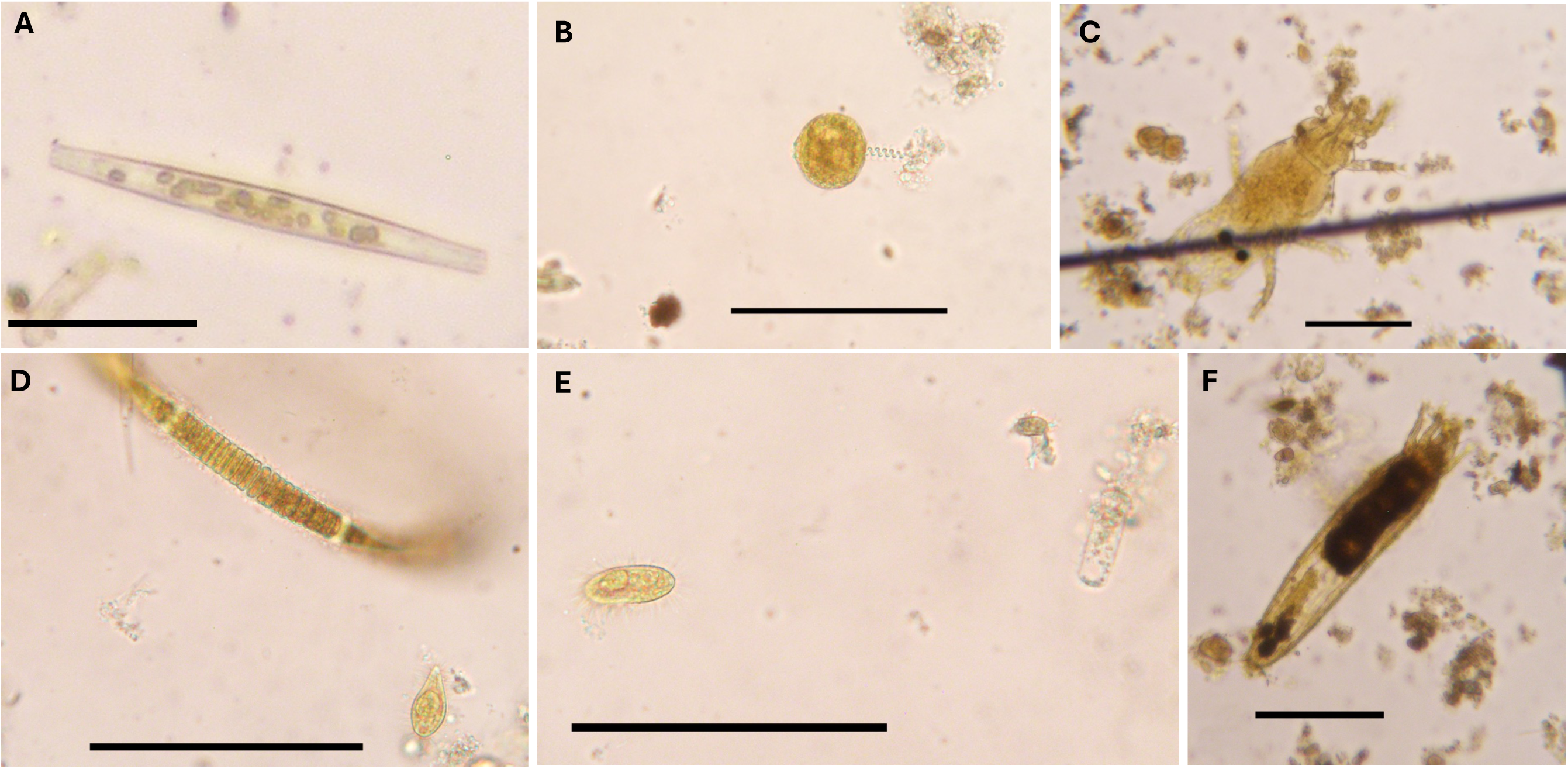
Microscopic images of slick SML biomass retrieved from the 100 µm mesh showing pennate diatoms (A), *Vorticella*-like ciliate (B), filamentous cyanobacteria accompanied by ciliate 1 (D), two ciliates (E) and two metazoans (C & F, respectively) The surroundings show excessive debris and decomposing matter. Scale bar represents scale at 100 µm.

## Discussion

Our study advances the understanding of the biogeochemical role of slicks by highlighting their impact on microbial distributions and NCP of O_2_ in the Baltic Sea. Our data show that abundances of eukaryotic pico- and nanophytoplankton as well as cyanobacteria are strongly correlated, suggesting tight community coupling of phototrophs of different size classes in the SML and surface enrichment in slicks of the Baltic Sea. These organisms are characterized by their small size, buoyancy, an apparent tolerance for high light conditions (Joint et al. 1990), and a preference for high nitrate supply (Otero-Ferrer et al. 2018) (Figure 3B). Cyanobacteria and eukaryotic picophytoplankton are often enriched in the SML, slicks, and associated foams (Rahlff et al. 2021; Rahlff et al. 2023b; Wang et al. 2014; Wurl et al. 2016). Diazotrophic cyanobacteria may dominate under low inorganic nitrogen conditions by fixing atmospheric N2, as suggested by their negative correlation with TDN. TDN was likely dominated by dissolved organic nitrogen (DON). Supporting this, data from pier sampling of ULW nearby Heiligendamm (Osterholz et al. 2021) on 18^th^ June 2024 showed low nitrate (0.2 µmol L⁻¹) and phosphate (0.03 µmol L⁻¹), indicating N limitation. Elevated silicate concentrations (9.2 µmol L⁻¹), compared to levels observed the week before and after (<5 µmol L⁻¹), suggest a transient summer bloom (data from personal communication with J. Waniek and J. Kuss). Cyanobacteria previously found in the Baltic Sea SML of the sampling region are *Nodularia spumigena*, *Anabaena* sp., and *Coelosphaerium* sp. (Stolle et al. 2011). SML prokaryotes correlated with NCP of O_2_ in the SML suggesting they play a role in carbon and O_2_ cycling. Based on previous findings, it is likely that only specific bacteria, such as Gammaproteobacteria, accumulate and thrive in the SML of slicks (Rahlff et al. 2023b). The ULW, that lies just below the SML, is less affected by direct atmospheric inputs. Microphytoplankton may sink below the SML, a process potentially enhanced by reduced surface tension due to surfactant accumulation (Sturdy and Fischer 1966). It can also be a strategy of microalgae to avoid surface stressors such as ultraviolet radiation and photoinhibition, which are common in the SML (Albright 1980; Hardy and Apts 1984). Prokaryotes showed positive correlations with microphytoplankton in the ULW, possibly benefiting from organic matter released by the latter and contributing to nutrient recycling. Microphytoplankton was negatively correlated with surface EFs of pico- and nanophytoplankton (N2), suggesting a vertical niche separation among phytoneuston and phytoplankton of different sizes. In addition, our microscopy data, even if available for only one sample, indicate that slicks can serve as reservoirs for juvenile stages of macrophytes. The high density of ciliates instead of presence of copepods and cladocerans could possibly indicate a food web that’s reliant on picophytoplankton and heterotrophic bacteria rather than larger microalgae. Nematodes, as a third trophic level preying on ciliates, add a step to the food web that may reduce energy transfer efficiency compared to copepods and cladocerans feeding directly on large microalgae (Pomeroy 1974). However, our methods did not capture the 50-100 µm size range.

Slicks are known to influence horizontal spreading dynamics (Hale and Mitchell 1997). However, the vertical spreading dynamics of neustonic organisms from the SML or slicks due to lowered surface tension have received little attention. Our data suggest that surfactants may play a role in structuring microbial communities across the air-sea interface, potentially by altering physicochemical properties of the SML or through ecological interactions such as inhibition or competition for resources. The observed negative correlation between surfactant concentrations in the SML and prokaryotic abundance in the ULW supports the idea that surfactants could influence microbial distribution and activity across layers. Passive sinking may be complemented by active microbial migration, as motility and chemotaxis genes have been detected in bacteria from a Baltic

Sea slick (Rahlff et al. 2023b). Aerotaxis, i.e., the directed movement in response to O_2_ concentration gradients, can influence the behavior of bacteria and algae by guiding them toward or away from oxygen-rich zones (Norkrans 1980; Roveillo et al. 2020), such as those found at the air-sea interface. Cyanobacteria, for example, regulate their buoyancy by means of intracellular gas vesicles (Walsby et al. 1995). Our results suggest that in the Baltic Sea, slick-induced enrichment of microorganisms in the SML and ULW may significantly influence oxygen dynamics, especially in increasingly hypoxic nearshore areas where reduced turbulence and enhanced stratification - exacerbated by warming - can amplify the impact of slicks on gas exchange (Andersen et al. 2017; Conley et al. 2011; Mustaffa et al. 2020). Our findings highlight the complexity of SML processes and the need for further studies to resolve fine-scale variability in oxygen gradients in relation to neustonic and planktonic microorganisms in and under naturally formed sea slicks.

## Acknowledgements

We acknowledge the Leibniz Institute for Baltic Sea Research, Warnemünde (IOW) and particularly Daniel Herlemann for hosting BCC and JR as guest scientists to conduct field work and sample processing. We gratefully acknowledge Heide Schulz-Vogt and Christin Laudan for providing access to laboratories at the IOW. We thank Madleen Dierken, Lars Kreuzer, and Birgit Sadkowiak for Heiligendamm pier nutrient data, and Jenny Jeschek for technical assistance. We further acknowledge excellent support by Danilo Erdmann and his team from the Sportbootverleih Warnemünde. BCC received funding from the German Research Foundation (DFG) grant

“NFDI4Microbiota” number NFDI 28/1 (DFG project number 460129525). JR received funding by the Swedish Research Council, Starting Grant ID 2023-03310_VR.

## Author contribution statement

Carolin Peter conducted microscopy analysis. Helge-Ansgar Giebel conducted cell count measurements and analysis. Bingli Clark Chai, Tassiana S. G. Serafim, Janina Rahlff conducted water sampling. Helena Osterholz was responsible for DOC measurements and analysis. Carola Lehners and Oliver Wurl were responsible for surfactant measurements and analysis. Janina Rahlff conducted oxygen measurements and analysis, correlations and together with Carolin Peter wrote the first draft. All authors contributed to manuscript editing.

## Scientific significance statement

The sea-surface microlayer (SML) is a dynamic boundary layer that regulates the exchange of gases, nutrients, and organic matter between the ocean and atmosphere. Slicks often form within this layer, concentrating microorganisms and altering physical and chemical conditions. While the SML is known to influence biogeochemical fluxes, the combined effects of microbial abundance, oxygen consumption, and surfactant presence are still poorly understood, especially in slicks of productive coastal systems. By examining oxygen consumption in relation to microbial counts, surfactants, and dissolved elements, we show that slicks enhance microbial respiration and drive vertical niche partitioning among plankton groups. These findings highlight the role of slicks as localized biogeochemical hotspots that modulate oxygen dynamics at the ocean-atmosphere interface.

## Data availability statement

The data are provided in the supplementary as material. Microscopy images can be obtained from Figshare doi: 10.6084/m9.figshare.29336453 (Peter and Rahlff 2025).

