## Supplement material for "Biogeochemical function of slicks in coastal surface waters of the Baltic Sea"

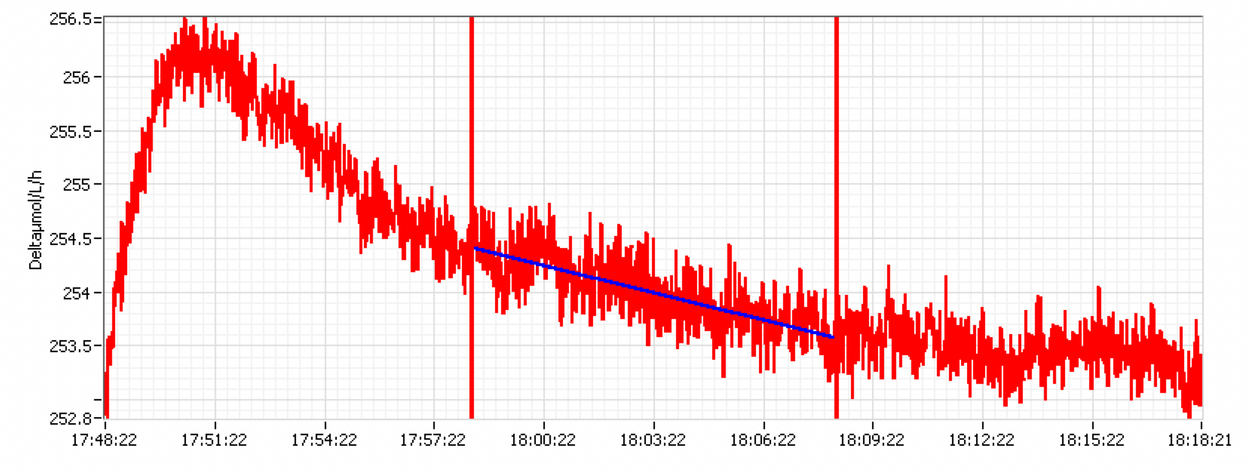


**Figure S1:** Representative figure for delta $O_2$ rate with the 10 min range for rate extraction.

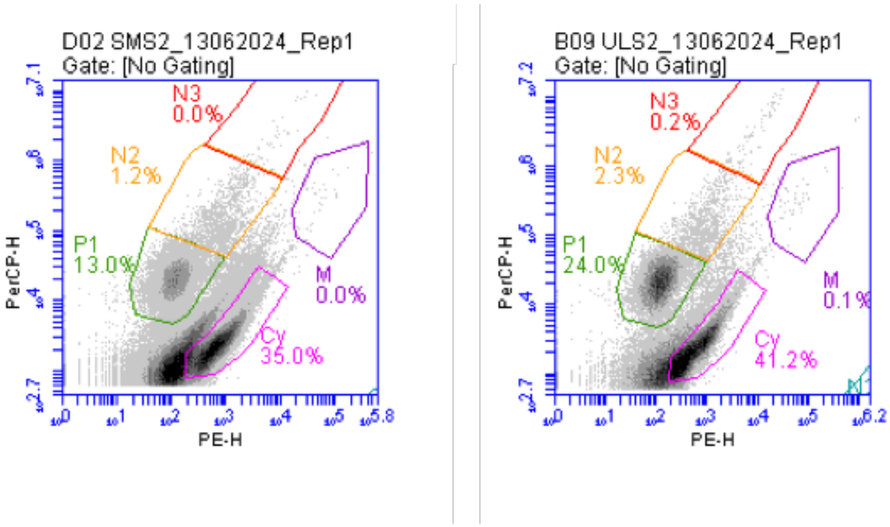


**Figure S2:** Flowcytometric gating of eukaryotic phototrophs on red (PerCP) vs. orange fluorescence (PE) after (John et al. 2022). Phytoplankton populations: P1=pico-sized, N2 and N3= nano-sized, M=micro-sized, Cy=cyanobacteria.

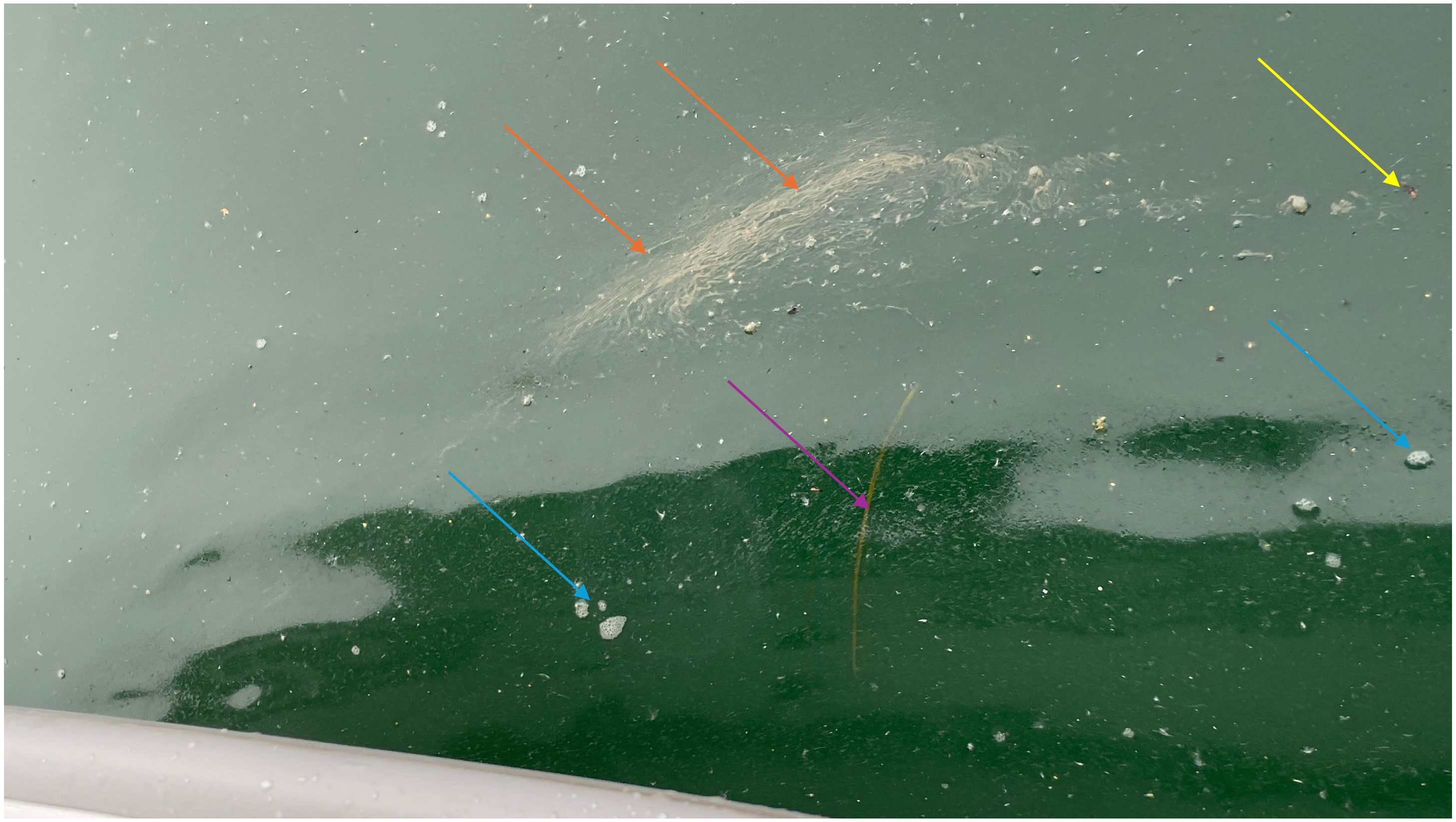

**Figure S3:** Viscous surface slick accumulating foams (blue arrows), insect (yellow arrows), seaweed (purple arrow) and chains of cyanobacteria (orange arrow) encountered at Day2, 16th June 2024 (photo: Tassiana Soares Gonçalves Serafim).

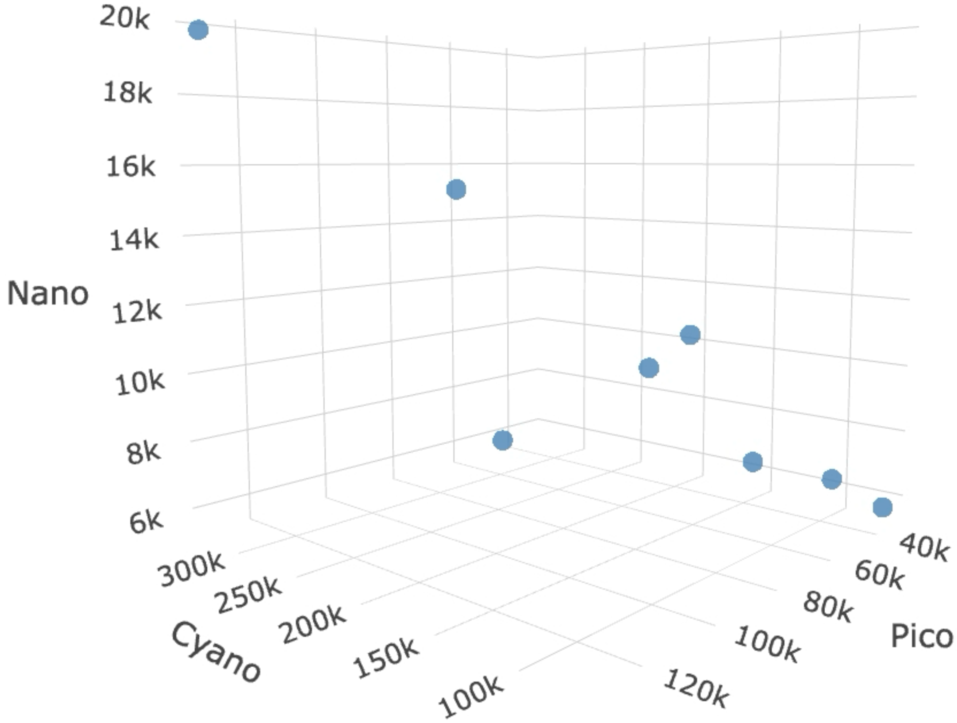


**Figure S4:** Correlation between cyanobacteria, pico and nano (N2) phototrophs in the SML.
